# A quantitative binding model for the Apl protein, the dual purpose recombination-directionality factor and lysis-lysogeny regulator of bacteriophage 186

**DOI:** 10.1101/784983

**Authors:** Erin Cutts, J. Barry Egan, Ian Dodd, Keith Shearwin

**Affiliations:** Department of Molecular and Biomedical Science, University of Adelaide, Adelaide, Australia, 5005; Division of Structural Biology, Institute of Cancer Research, London, United Kingdom

## Abstract

The Apl protein of bacteriophage 186 functions both as an excisionase and as a transcriptional regulator; binding to the phage attachment site (*att*), and also between the major early phage promoters (pR-pL). Like other recombination directionality factors (RDFs), Apl binding sites are direct repeats spaced one DNA helix turn apart. Here, we use *in vitro* binding studies with purified Apl and pR-pL DNA to show that Apl binds to multiple sites with high cooperativity, bends the DNA, and spreads from specific binding sites into adjacent non-specific DNA; features that are shared with other RDFs. By analysing Apl’s repression of pR and pL, and the effect of operator mutants *in vivo* with a simple mathematical model, we were able to extract estimates of binding energies for single specific and non-specific sites and for Apl cooperativity, revealing that Apl monomers bind to DNA with low sequence specificity but with strong cooperativity between immediate neighbours. This model fit was then independently validated with *in vitro* data. The model we employed here is a simple but powerful tool that enabled better understanding of the balance between binding affinity and cooperativity required for RDF function. A modelling approach such as this is broadly applicable to other systems.

## INTRODUCTION

Apl is one of a large family of recombination directionality factors (RDFs) (Lewis and Hatfull, 2001), that modulate the directionality of site-specific recombination reactions catalysed by the tyrosine integrase/recombinase proteins (Esposito and Scocca, 1997). RDFs are small proteins that bind to the DNA flanking the recombination site and, by altering the DNA architecture or by interacting with the integrase protein, act to alter the binding of the integrase complex to promote excision of the prophage.

RDFs appear to share a mode of DNA binding in which protomers bind with high cooperativity in a head-to-tail manner to tandem DNA repeats spaced one DNA turn apart, shown for λ Xis (Sam *et al.*, 2002), Gifsy-1 Xis (Flanigan and Gardner, 2007) and Pukovnik Xis (Singh *et al.*, 2014). The crystal structure of P2 Cox has been solved in the absence of DNA, revealing an extensive interaction with neighbouring cox protomers (i+1) and also interactions with i+2 (Berntsson *et al.*, 2014). Where examined, RDFs have been shown to cause large bends in attachment site (*att*) DNA (λ Xis (Thompson and Landy, 1988) (Cho, Gumport and Gardner, 2002), P2 Cox (Ahlgren-Berg *et al.*, 2009), L5 Xis (Lewis and Hatfull, 2003), P4 Vis (Calì *et al.*, 2004), Wϕ Cox (Ahlgren-Berg *et al.*, 2009), P22 Xis (Mattis, Gumport and Gardner, 2008) and Pukovnik Xis (Singh *et al.*, 2014)). The crystal structure of the archetypal RDF, Xis from λ, showed three Xis monomers bound to the X1-X1.5-X2 sites in *att*R causing a 72° non-planar bend in the DNA, leading to the hypothesis that a twisted microfilament forms (Abbani *et al.*, 2007), a hypothesis supported by DNA compaction studies on P2 Cox (Frykholm *et al.*, 2016). Another apparent common feature of RDFs is relaxed DNA specificity, with binding at non-canonical DNA sites seen *in vitro* at higher RDF concentrations. The crystal structure of lambda Xis-DNA complex also showed fewer sequence-specific contacts are made to the X1.5 site, compared with the X1 and X2 sites (Abbani *et al.*, 2007). Based on amino acid sequence, Apl is an outlier in the RDF family (Lewis and Hatfull, 2001) but appears to fit this DNA binding pattern, being monomeric in solution (Shearwin and Egan, 1996) and binding to the 186 *att* site, along with the integrase protein (Int) and the integration host factor (IHF), to five direct repeats sequences with 10-11 bp spacing (Dodd, Reed and Egan, 1993) (Fig. 1).

**Figure 1.**
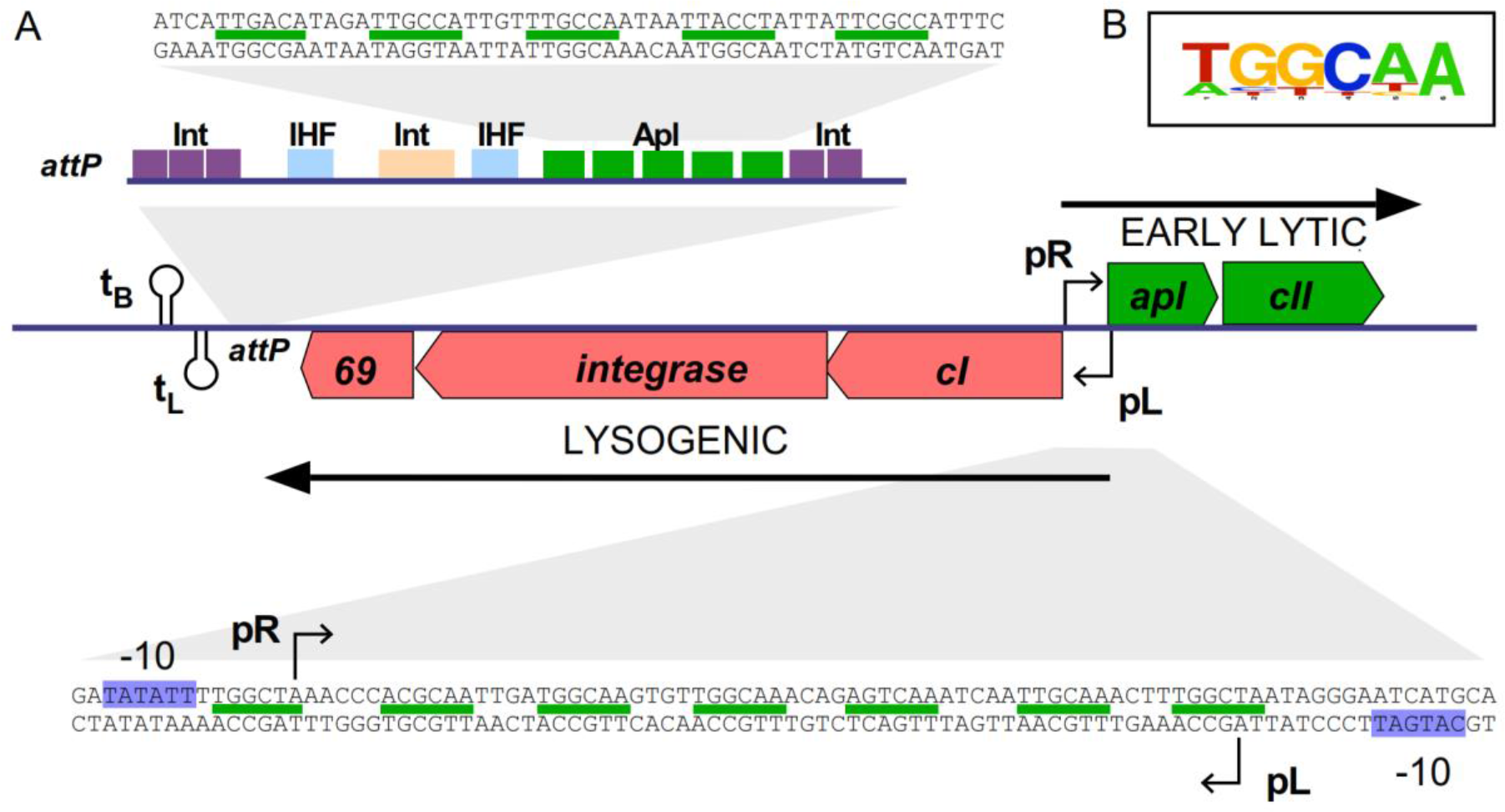
(A) Map indicating the location of Apl binding sites in bacteriophage 186. The centre of the diagram shows the switch region of 186, where the lytic pR promoter and the lysogenic pL promoter are arranged in a convergent orientation. There are seven Apl operators (indicated by green lines) located between the pR and pL promoters, and five Apl operators located at *att*P. Individual Apl operator sequences are six base pairs long, present in a direct repeat arrangement. At *att*P, the IHF sites are shown in blue and Integrase binding sites shown in tan (core site) and purple (arm sites). (B) Logo plot (Crooks *et al.*, 2004) representing the consensus of all Apl operator sequences from coliphage 186.

A subset of RDFs also function as transcriptional regulators. In the KlpE (Panis, Méjean and Ansaldi, 2007) and P4 (Calì *et al.*, 2004) prophages, the promoter for the integrase gene lies near the *att* site and is repressed by binding of the RDF to its sites within *att*. Such regulation is potentially widespread given the common proximity of *att* sites and *int* genes. Apl and other RDFs from P2-like bacteriophages and P4 also regulate transcription at locations well away from their attachment sites. These HTH-motif proteins are each encoded by the first gene of the phage early lytic operon and regulate the balance between lytic and lysogenic transcription, using recognition sequences of similar sequence and arrangement to the *att* sequences (Reed *et al.*, 1997)(Yu and Haggard-Ljungquist, 1993)(Esposito, Wilson and Scocca, 1997). In the P2-like phages, these RDF binding sequences typically lie between and overlapping the lytic and lysogenic promoters, which are arranged face-to-face and separated by 40-60 bp (Karlsson *et al.*, 2006)(Nilsson *et al.*, 2011). In 186, the region between the pR lytic promoter and the pL lysogenic promoter contains seven Apl recognition sequences that, like the sites at *att*P, are direct repeats with 10-11 bp spacing (Fig. 1). Apl binding represses both promoters (Dodd, Reed and Egan, 1993); however, while Apl has a clear function at the *att* site as an excisionase (Reed *et al.*, 1997), the function of its repressive activity at pR and pL is not well understood.

To better understand the mechanism of action and function of Apl, particularly with regard to its regulation of lytic and lysogenic transcription, we further investigated its mode of DNA binding. We show that purified Apl binds at pR-pL with high cooperativity, bends the DNA and spreads from specific binding sites into adjacent non-specific DNA. Although we were unable to detect Apl binding to a single site, we were able to use a simple mathematical model to extract estimates of binding energies for specific and non-specific sites and cooperativity by measuring Apl binding *in vitro* and *in vivo* to different numbers and arrangements of DNA sites. Each Apl monomer binds to DNA with low sequence specificity but with strong cooperativity between immediate neighbouring monomers.

## MATERIALS AND METHODS

### Assay and Expression Strains

NK7049 (ΔlacIZYA) X74 galOP308 Str^R^ Su^−^ from R. Simons (Simons, Houman and Kleckner, 1987) was the host strain for all LacZ assays. DH5α and XLI-blue were hosts for recombinant DNA work. Strains were grown at 37°C in lysogeny broth (LB), with the addition of ampicillin (100 μg ml^−1^ for pZE15 based plasmids) and kanamycin (50 μg ml ml^−1^ for pUHA1) where necessary.

pZE15-P_lac_-LacZ was constructed by inserting the *lacZ* gene into the *Bam*HI and *Hind*-III sites of the ampicillin resistant, colE1 based plasmid pZE15 (Dodd *et al.*, 2001). Lac repressor was supplied by pUHA-I, a p15A based plasmid encoding kanamycin resistance and carrying the wild-type lacI gene and promoter, obtained from H. Bujard (Heidelberg University, Germany).

Chromosomally integrated LacZ reporters were NK7049 (λRS45ΔYA pBC2-based or pMMR9-R) based 186 pR- or 186 pL-lacZ reporters. The pR- and pL-lacZ reporter plasmids were created as described in (Dodd and Egan, 2002). These plasmid-based lacZ fusions were then transferred to the lacZ reporter phage λRS45ΔYA for insertion into the *E. coli* chromosome. Plasmid-containing strains were infected with λRS45ΔYA, and blue plaque-forming phage among the progeny were identified and purified on NK7049, on plates containing X-gal (Simons, Houman and Kleckner, 1987). Lysogenising NK4079 with the reporter phage ensures that the reporters are all located at an identical position (*att* lambda) in the chromosome. Chromosomal integrants were checked for monolysogens by PCR (St-Pierre *et al.*, 2013). Apl, or mutants thereof, were supplied to the reporter strains by pZE15Apl, a colE1 based plasmid (Dodd *et al.*, 2001), where Apl expression was under control of the plac promoter. Reporter strains also carried the pUHA-1 plasmid, as a source of lac repressor. Thus, Apl expression was controlled by addition of IPTG to the growing culture, and promoter activities assayed in a microtiter plate format, according to Palmer *et al.* (Palmer *et al.*, 2009).

Expression and purification of recombinant Apl protein from BL21 pLysS pET-Apl was performed as previously described (Shearwin and Egan, 1996). For experiments with mutated Apl, Apl expression levels were assessed by Western blots, using a method described previously (Cui *et al.*, 2013). The anti-Apl polyclonal antibody was generated in a rabbit against the KLH-conjugated Apl peptide ASEIAIIKVPAPIVC by Genscript (New Jersey, USA).

### *In vitro* DNA Binding Assays

#### Gel mobility shift assays

For gel shift assays, double stranded DNA fragments with one strand ^32^P end-labelled were generated by PCR in which one of the primers had been ^32^P end-labelled using polynucleotide kinase. The double stranded ^32^P-labelled PCR product was purified by polyacrylamide gel electrophoresis, the DNA eluted from the gel slice overnight at 37°C, ethanol precipitated and resuspended in binding buffer before use.

The DNA sequences of the oligonucleotides used in binding assays are given in Supplementary Fig. 1.

Binding reactions (10 μL) were prepared by addition of DNA (∼5 cpm), Apl (exhaustively dialysed against 50 mM Tris-HCl (pH 7.5), 0.1 mM EDTA, 10% (v/v) glycerol, 150mM NaCl (TEG 150)) and binding buffer (TEG150). Reactions were left on ice for at least 30 minutes to allow attainment of equilibrium, and 6 μL loaded onto running polyacrylamide (0.5 X TBE) gels containing 10% glycerol. For binding to short DNA fragments, 15% gels were used, while 8% gels were used for DNA bending assays. Gels were electrophoresed at 4°C at constant current (20mA) for approximately 2 hours. Upon completion of electrophoresis, gels were dried, exposed to a phosphorimager screen and quantitated using the volume integration feature of Imagequant (Molecular Dynamics) or Imagelab (BioRad) software.

The fraction of DNA bound in each lane was calculated as (counts for the retarded band) / (counts for the whole lane), and corrected for a small degree of protein independent smearing using a no protein control lane. The DNA concentration was sufficiently low that total protein concentration could be substituted for free protein concentration.

#### Bending assay

DNA fragments containing 3, 4, 5, 6 or 7 Apl binding sites (Supplementary Fig. 1) were prepared by annealing complementary oligonucleotides and ligating into the blunt *Hpa* I site of pBend 5 (Zwieb and Adhya, 2009). The region of this plasmid containing 17 circularly permuted restriction sites and the Apl binding sites were then amplified by PCR using primers pBend SK (TAGTGGATCCCCCGGGCTGCA) and pBend KS (CGACGGTATCGATAAGCTTGG). This fragment was labelled by inclusion of ^32^P α-ATP (10 μCi) in the PCR reaction. The PCR product was purified by polyacrylamide gel electrophoresis and an aliquot digested in 10 μL reactions with *Mlu*I, *Eco*RV or *Bam*HI. Digestion produced three fragments, including a fragment containing the Apl binding site located at either the left end, centre or right end within the fragment. Two μL of this digest was used in binding reactions for determination by gel shift assay of the electrophoretic mobility of the protein-DNA complex. Binding reactions were performed in TEG 50 buffer and contained 3.2 μM Apl. Samples were loaded on 8% polyacrylamide, 0.5X TBE gels containing 10% glycerol and run at 20 mA constant current and 4°C. Loading dye was run in a separate lane. Following electrophoresis, gels were dried, exposed to a phosphorimager screen, and DNA mobility quantitated using Imagequant software (Molecular Dynamics). The apparent bend angles were quantitated according to equation 1.

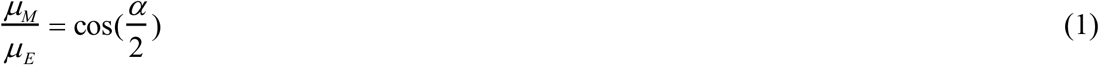

where α is the bend angle and *μ*_*M*_ and *μ*_*E*_ are the relative mobility of DNA fragments containing the binding site at the middle and at the end of the fragment, respectively (Zwieb and Adhya, 2009). Apparent bend angles were calculated from the mean of four independent experiments.

#### DNAseI footprinting

Experiments were performed essentially according to (Sandaltzopoulos and Becker, 1994), with modifications described by (Shearwin and Egan, 2000). This method uses magnetic beads to facilitate sample preparation. Double stranded DNA fragments for footprinting were prepared by PCR using a ^32^P end-labelled primer and a biotinylated unlabelled primer (biotin-RSP). The PCR reaction (20 μL) was passed over a PCR purification spin column (Geneworks, Adelaide) to remove any unincorporated biotinylated primer which would compete with full length product for binding to the beads. The eluate from the spin column (60 μL) was added to 75 μL of streptavidin-coated magnetic beads (Dynabeads, Dynal), prepared according to the manufactures recommendations, and incubated for one hour at room temperature to allow the biotinylated, radiolabelled PCR product to bind. The beads were then washed several times, resuspended in 50 - 100 μL binding buffer and stored at 0 °C for up to 1 week. Bead DNA (5 μL, ∼6000 cpm) was added to binding buffer containing appropriate Apl concentrations, in a total volume of 40 μL. The footprint binding buffer consisted of 50 mM Tris-HCl (pH 7.5), 0.1 mM EDTA, 10% (v/v) glycerol, 75 mM NaCl, 10mM MgCl_2_ 1.5 mM CaCl_2_ 1 μM bovine serum albumin (BSA). These binding reactions were incubated at 37 °C for at least thirty minutes to allow attainment of equilibrium, prior to addition of DNase 1 (0.5 ng). The DNase 1 reaction was allowed to proceed for exactly ten minutes at 37 °C before being stopped with 50 μL of stop solution (4M NaCl, 100 mM EDTA). The beads were washed once with 100 μL of 2M NaCl, 20 mM EDTA, once with 100 μL of 10 mM Tris-Cl, 1 mM EDTA, pH 8.0 and resuspended in 6 μL of loading buffer (90% formamide, 10 mM EDTA). The reactions were heated to 90 °C for 3 minutes and 5 μL loaded immediately onto a 6% denaturing polyacrylamide gel. Electrophoresis was at 1500 V (constant voltage) for approximately 2 hours. Following electrophoresis, gels were dried onto filter paper, exposed overnight to a phosphorimager screen and viewed using Imagequant. Apl concentrations used in the footprints were: 3000, 2000, 1000, 794, 631, 500, 400, 319, 100, 10 nM.

### *In vivo* Apl expression system

Apl expression from pZE15Apl was controlled from the pLac promoter by Lac repressor supplied by pUHA-1. Relative expression of Apl from pZE15Apl in NK7049 λRS45ΔYA-pMRR9R-MMpR^+^pL^+^.lacZ from the pLac promoter at 0, 3.6, 5.4, 7.2, 9, 13.5, 18, 20, 27, 40, 60, 100 μM IPTG was determined by comparison with lacZ expression from pZE15-plac-LacZ in NK7049.

Data was pooled from assays performed on two to three different days, each with 4 biological replicates. For each data set, the Apl-containing lacZ reporter value was divided by the mean parental (no Apl) plasmid value for that IPTG concentration, and relative repression pooled. The 95% confidence intervals of the pooled data were calculated, and relative repression curves plotted with y-axis error bars being 95% confidence intervals in relative repression value and x-axis error bars being 95% confidence intervals in pLac promoter activity.

### Statistical Mechanical Modelling

A full description of the statistical mechanical model and optimisation of parameter fitting is given in Supplementary Materials.

## RESULTS

### Three adjacent operators are required for efficient Apl binding *in vitro*

Purified, refolded Apl protein (Shearwin and Egan, 1996) was used in gel shift assays to determine the number and arrangement of recognition sequences needed for efficient binding *in vitro* (Fig. 2). Binding was detected only when three or more operators were present, suggesting that Apl binding is highly cooperative. The apparent K_D_ decreased as the number of operators was increased. When six or seven operators were present, multiple shifted species were observed, demonstrating multiple, distinct relatively stable Apl-DNA complexes.

**Figure 2.**
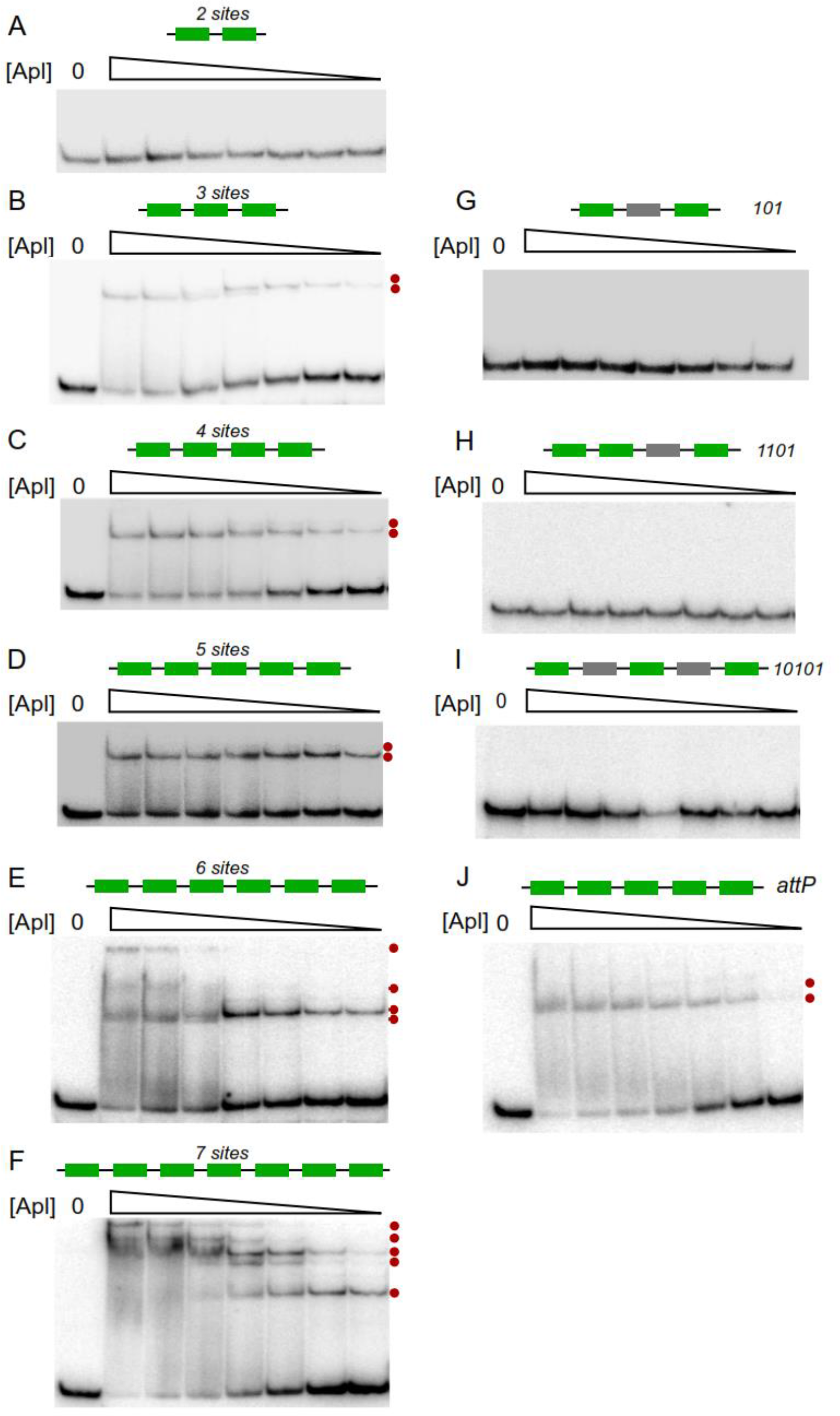
Apl DNA binding to DNA. DNA gel shift assays using DNA oligomers containing three to seven Apl (A-F) binding sites. Binding was assayed to DNA fragments with various combinations of specific and non-specific sites; 101 (G), 1101 (H) and 10101 (I), where 1 represents a specific Apl operator and 0 represents a scrambled operator. An assay was also performed on a fragment containing the 5 specific operators from the *att*P site (J). Apl concentrations were 6400, 3200, 1600, 800, 400, 200, and 100 nM. Red dots next to a gel indicate the presence of a different protein-DNA species.

To examine whether cooperation between multiple operators is adjacent or longer range, binding was tested using DNAs with combinations of intact and mutated Apl recognition sequences. These were designated 101, 1101 and 10101, where 1 indicates an intact operator and 0 indicates a scrambled site. No binding was detected to any of these fragments (Fig. 2G-I). The binding to the 111 fragment, but lack of binding to the fragment containing scrambled sites, indicates that three adjacent operators are needed for efficient binding *in vitro*. A gel shift experiment using the five operators at *att*P showed a similar binding pattern to that seen using the central five operators at pR-pL, indicating a similar mechanism of binding.

### Apl bends DNA upon binding

As DNA bending appears to be a conserved property of the tyrosine integrase family of RDF proteins, the ability of Apl to bend DNA was tested more directly using the ‘circularly permuted gel shift’ technique (Fig. 3) (Kim *et al.*, 1989)(Zwieb and Adhya, 2009). Fragments containing three, four, five, six or seven adjacent operators from pR-pL were tested. Retardation of the fragments differed markedly depending on the position of the set of binding sites within the fragment, suggestive of bending. The calculated bend angle (from Equation 1) for the 3-operator segment was 87±1° (Fig. 3). This value is similar to the 72° seen in the crystal structure of λ Xis bound to three adjacent sites (Abbani *et al.*, 2007), though smaller than the 120° estimate using a similar gel shift technique (Thompson and Landy, 1988). Increased apparent bend angles were obtained for the four-, five- and six-operator segments, with a possible slight decrease for the seven-operator segment (Fig. 3). These bends are of similar magnitude to those seen using the same technique with P22 Xis (Mattis, Gumport and Gardner, 2008), P2 Cox (Ahlgren-Berg *et al.*, 2009) and Puckovnik Xis (Singh *et al.*, 2014) at their attachment sites. As the angle estimate is based on an assumption of a planar bend, which may not be the case for DNA with multiple Apl operators, these changes in apparent bending indicate only that the architecture of the DNA changes with increasing bound Apl.

**Figure 3.**
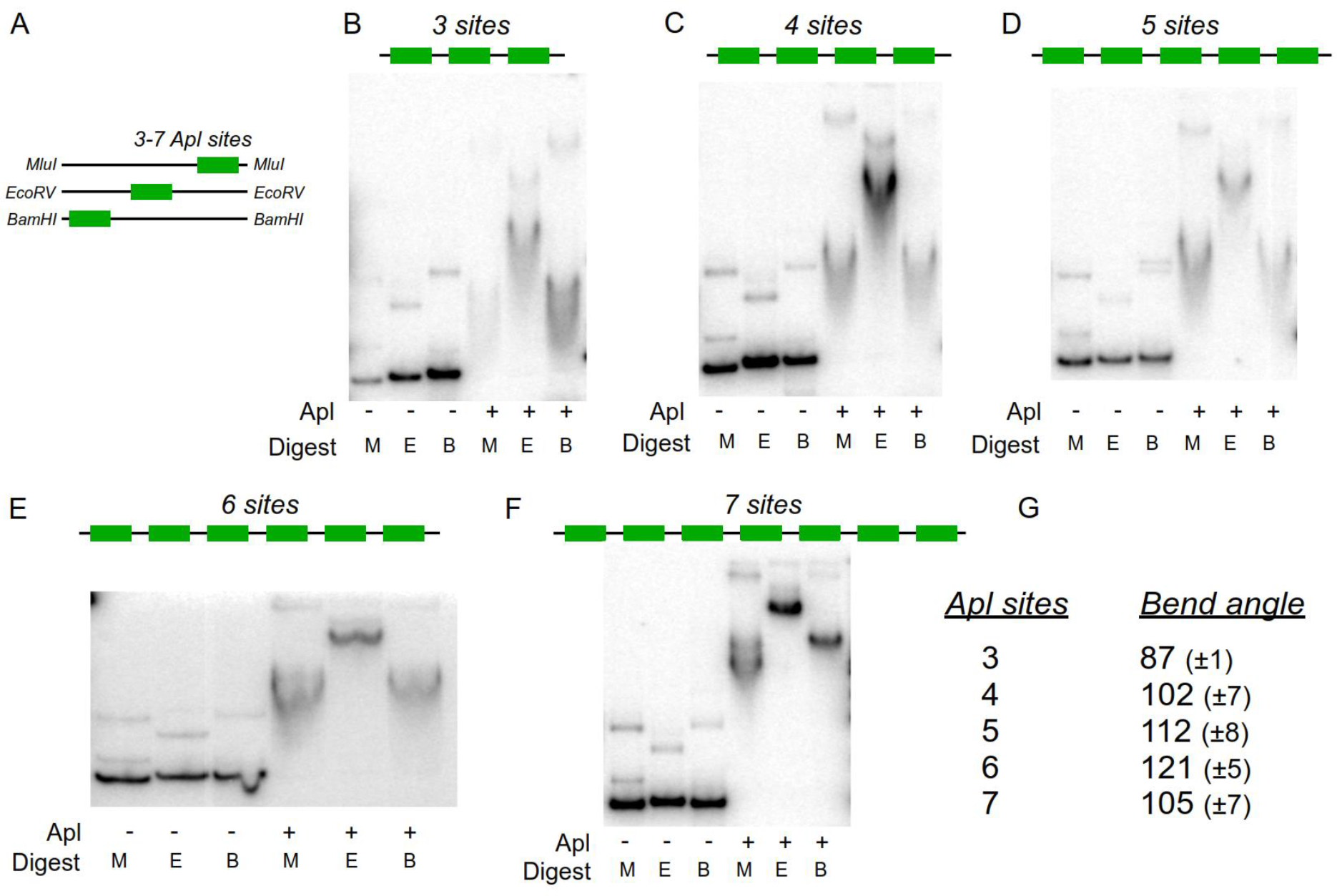
DNA bending assay. (A) The arrangement of Apl binding sites at the ends or centre of the DNA fragment allows estimation of DNA bending angle from gel shift assays. (B-F) shows the gel shifts in the absence and presence of 3.2 μM Apl for fragments containing 3-7 Apl operators. M, E, B indicate the fragment corresponding to digestion with *Mlu*I, *Eco*RV or *Bam*HI, as in panel A. (G) Summary of apparent bend angles derived from the bending assays. Errors are confidence limits based on four independent experiments.

### Apl binding spreads into adjacent non-specific sites

While Apl DNase I footprinting performed by (Dodd, Reed and Egan, 1993) revealed protections and enhancements that extended beyond the specific binding sites at pR-pL and at *att*P, these assays were performed with crude cell extracts and with native 186 sequences flanking the operator DNA. Hence, the observed DNA alterations may have been due to proteins other than Apl or to Apl binding to cryptic operators within the 186 sequence. To test for spreading of Apl binding into adjacent non-specific sequences, DNAse I footprinting was repeated using purified Apl and with either three (Fig. 4A) or five (Fig. 4B) adjacent Apl operators embedded in non-186 DNA. In both cases, the periodic protections and enhancements previously observed within the Apl operators (Dodd, Reed and Egan, 1993) were apparent. Furthermore, protections continued on either side of the specific sequences, showing Apl spreading to non-specific sites in the vector DNA, with the extent of spreading increasing with Apl concentration.

**Figure 4.**
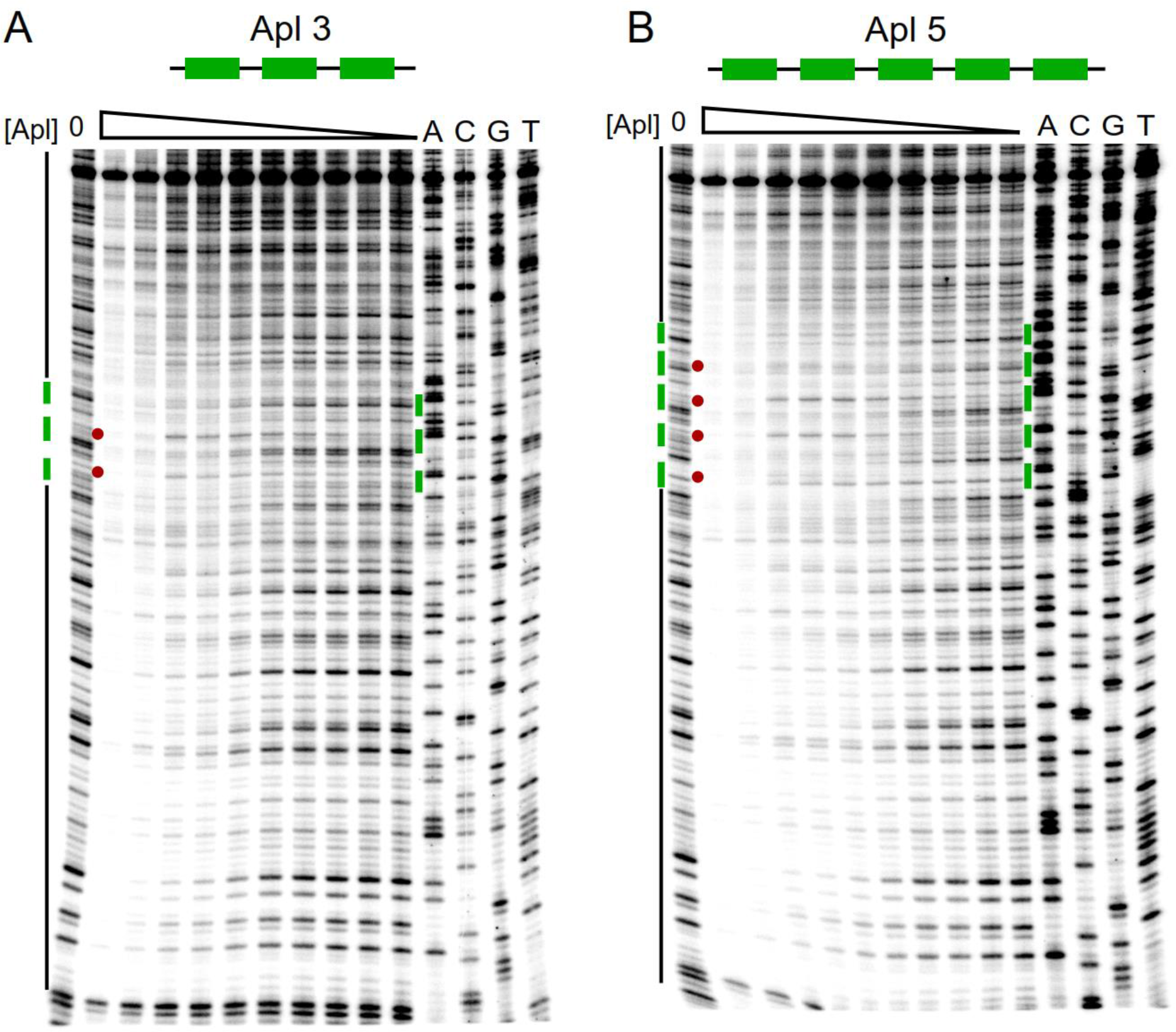
DNAse I footprinting shows spreading of Apl binding into surrounding DNA of unrelated sequence. DNA containing either (A) five or (B) three Apl operator sequences was embedded into unrelated DNA sequence (pBluescript plasmid). DNaseI footprint reactions, examining the equivalent of the 186 top strand, were performed as described in methods. The leftmost lane in each gel contained no Apl. Apl concentrations were 3000, 2000, 1000, 794, 631, 500, 400, 319, 100, 10 nM (left to right). The four lanes on the right hand side of each gel contain dideoxy sequencing reactions as indicated. At the side of each gel the green bars indicate the region of DNA corresponding to Apl operator sequences, and the black lines indicate sequence corresponding to plasmid DNA. The red dots represent the characteristic enhancements of DNAse I cleavage seen with Apl binding, indicative of distortion of the DNA (Dodd et al., 1993).

Interestingly, there is no sharp transition between occupation of the specific operator sites and occupation of the adjacent non-specific sites. That is, protection of the flanking DNA begins at Apl concentrations at which the operator sites are not fully occupied. This suggests that Apl’s affinity for its operators is not substantially greater than for non-specific sites. If these affinities were very different, then one would have expected that some Apl concentrations would give strong protection of the operator sites with very little spreading.

These results show that, like other RDFs, Apl binding at specific sites can seed spreading of Apl into adjacent non-specific sites, and indicate strong binding cooperativity and weak discrimination between the operator sites and non-specific DNA.

### Effect of operator mutations on Apl repression of the pR and pL promoters *in vivo*

To examine the role of cooperative binding of Apl on its activity at pR-pL, we used lacZ reporter constructs carrying mutations at the central three Apl operators. Three different fragments were tested: 1111111 (wild-type (WT), bearing operators 1-7), 1101011 (operators 3 and 5 scrambled) and 1100011 (operators 3, 4 and 5 scrambled). Fragments oriented to report either pR activity or pL activity were fused to lacZ in a lambda prophage, as described in Methods. The pL reporter carried mutations inactivating pR, in order to remove pR’s inhibition of pL by transcriptional interference (Callen, Shearwin and Egan, 2004). There is no reciprocal interference of pL on pR (Callen, Shearwin and Egan, 2004). Apl was supplied from a multicopy plasmid (pZE15Apl) under plac/IPTG control. We have not measured absolute Apl concentrations resulting from this expression system, however relative Apl expression was quantitated by constructing an equivalent pZE15-plac.lacZ plasmid and assaying LacZ activity in response to IPTG (Supplementary Fig. 2).Thus, relative Apl concentrations are given in terms of expression units.

Induction of Apl expression with IPTG resulted in repression of both pR and pL (Fig. 5A). Promoter expression is expressed relative to the Apl minus control plasmid (∼800 units for pR, ∼100 units for pL(pR^−^)). Wild type pR was repressed by Apl to ∼0.4 of unrepressed and wild-type pL(pR^−^) to ∼0.2, a difference in sensitivity consistent with previous observations (Reed *et al.*, 1997). Testing the effect of the 11001011 and 1100011 operator mutations on pR repression showed that in both mutants repression was much weaker but was still detectable at the highest Apl concentrations (Fig. 5A).

**Figure 5.**
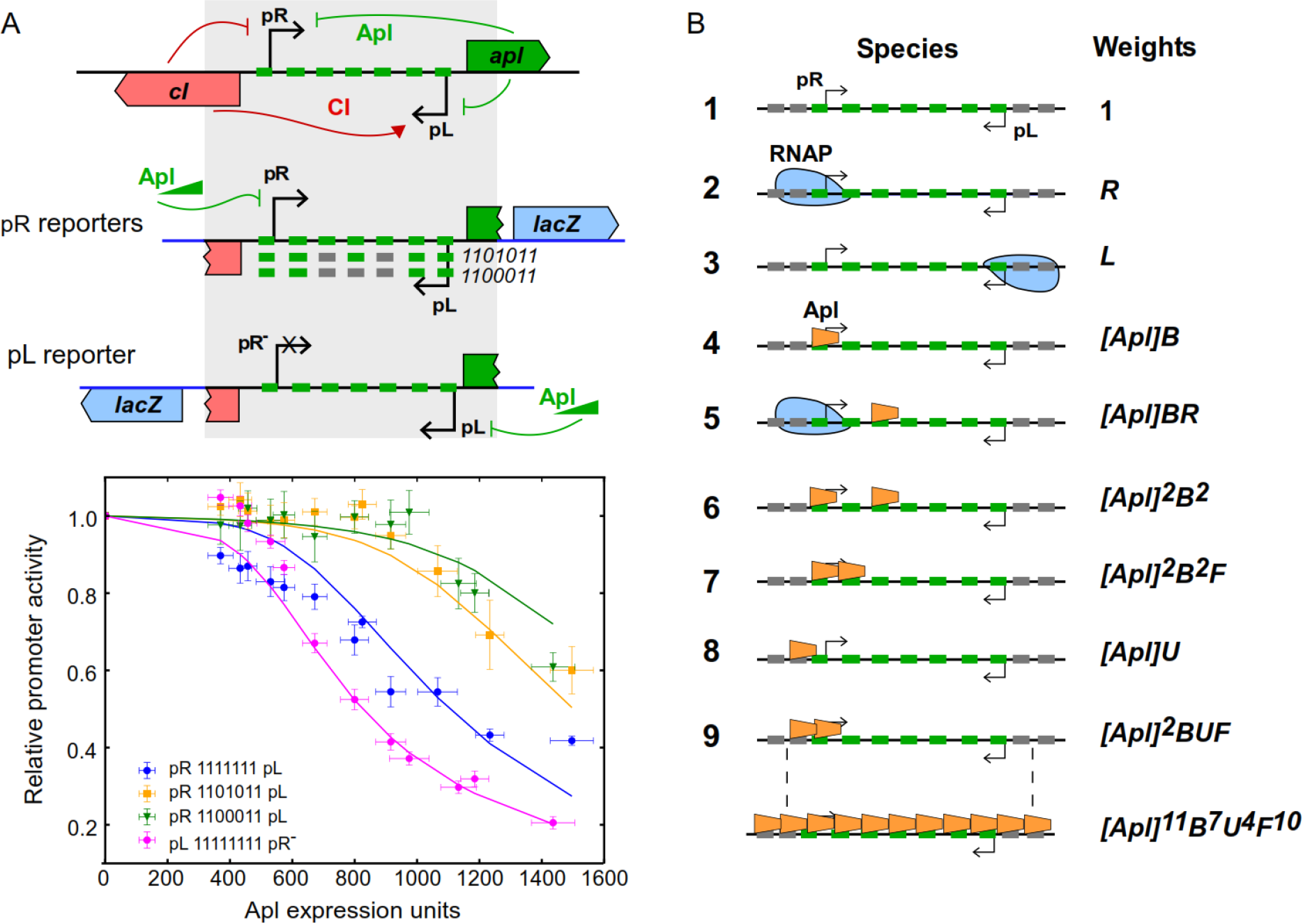
Mechanistic modelling of Apl DNA binding. (A) Explanation of model parameters, R and L for RNAP binding to pR and pL respectively, B and U, for specific and non-specific DNA binding respectively, scaled by Apl concentration. When two Apl monomers are bound adjacently, this is further multiplied by the cooperativity parameter, F. More complex states are described by multiplication of these parameters. (B) Top: the LacZ reporter measures Apl’s function as a repressor of the switch region, where it represses both pR and pL. Middle: pR drives expression of a LacZ reporter gene, this construct was also made with operators 3 and 5, and 3, 4 and 5 scrambled. Bottom: pL drives the expression of LacZ, with the pR promoter mutated to prevent transcriptional interference. (C) LacZ reporter data, with fit shown as solid lines. Error bars indicate 95% confidence intervals.

### Modelling of Apl repression of pR and pL

To achieve a more quantitative understanding of the mechanism of Apl regulation of pR and pL, we tested whether a simple statistical mechanical model of Apl DNA binding could explain the reporter data and quantify cooperativity and DNA binding affinity. The pR-pL region was modelled as comprising seven specific Apl operators (O_1_-O_7_), two non-specific Apl binding sites on each side (N_−1_, N_0_, N_8_ and N_9_) and two RNAP binding sites (Fig. 5B). Apl binding at operator site O_1_ and O_2_ was assumed to compete with RNAP binding at pR, since the conserved sequences lies at pR −1 to −6, and +5 to +10, respectively, as was binding to the two non-specific sites N_−1_ and N_0_ adjacent to site 1. The conserved 6 bp O_7_ sequence lies at pL +3 to +8 and Apl binding to this site or the adjacent non-specific N_8_ and N_9_ sites was also assumed to compete with RNAP binding to pL (Fig. 1).

All possible states were then defined and given a statistical weight based on the interactions present (Fig. 5B). Binding of an Apl monomer to an operator was given a statistical weight [Apl].B, where B is the specific association constant, and [Apl] is a scaled Apl concentration. Non-specific Apl binding of an Apl monomer was given a weight [Apl].U, where U is the non-specific association constant. A cooperation parameter, F, was applied for two adjacent bound Apl monomers. RNAP binding to the pR and pL promoters was given weights R and L, respectively, combining the unknown but constant cellular concentration of RNAP and unknown RNAP binding constants to the promoters. Based on these parameters, a weight for each of the 2448 possible states can be calculated. The probability of any state occurring is the weight of that state divided by the sum of the weights for all states. The activity of each promoter was assumed to be proportional to the sum of the probabilities of all the states where the RNAP is bound to the promoter. Thus, possible specific effects of Apl on transcription initiation, promoter clearance or elongation were ignored. A scaling factor for each promoter was applied to set its activity in the absence of Apl to 1.

The model parameters were adjusted to optimize the fit to the complete set of *in vivo* repression data, using a combined Monte-Carlo/linear optimization approach (Supplementary Methods; Supplementary Figs. 3 and 4). The repression of the WT pL(pR^−^) and WT pR reporters by Apl are reproduced well at higher [Apl] concentrations, however, the model predicts a greater difference between the 1101011 and 1100011 reporters than is seen in the data (Fig. 5A). The obtained Apl binding constants were B=4.55 (±0.53) ×10^−5^ and U=0.99 (±0.09) ×10^−5^ (Apl expression units)^−1^. Thus, the K_D_ for monomer binding to a single operator is ∼22, 000 Apl expression units (1/B). Since the maximum *in vivo* Apl concentration, produced from pLac on a multicopy plasmid, was ∼15-fold less than this (1500 expression units; Supplementary Fig. 2), it is clear that binding to a single operator is weak. The RNAP binding values were fit as R=6.88 (±1.09), and L=0.29 (±0.15) and although these values are harder to interpret as they are used to normalise the binding curves, the fact R is larger than L is consistent with pR being a stronger promoter.

Remarkably, non-specific binding is predicted to be only ∼4.5-fold weaker than specific binding. The fitted value for cooperativity, F=50.7 (±7.9), equivalent to a ΔG of −2.4 kcal/mol, reflects a large contribution to Apl binding from cooperativity between adjacent monomers. The apparent K_D_ for cooperative binding to two operators becomes ∼3100 expression units, only twice the maximum expression level, while for three operators the apparent K_D_ is ∼1600 expression units. For seven consecutive operators, the apparent K_D_ falls to ∼750 expression units.

We explored more complex versions of the model, such as having different site binding strength according to sequence variation and adding in an additional cooperation term as the P2 Cox structure suggests an i+2 contact (Berntsson *et al.*, 2014), but these models did not converge to one unique solution and did not significantly improve the data fit. Although the parameter values obtained with the more complex models were often different, the basic observations of high cooperativity and low discrimination were robust. Thus, a simple model of Apl binding applied to the *in vivo* repression data was able to confirm the qualitative conclusions from the gel shift and footprinting experiments that Apl binds with high cooperativity and with low discrimination between specific and non-specific sites.

### Modelling *in vitro* Apl-DNA binding

To test whether the Apl binding model is consistent with the *in vitro* Apl binding data, we tested whether it could reproduce key features of both the gel shift and DNAseI footprinting data. To obtain binding constants in units of Apl concentration, rather than Apl expression units, the value of B was fitted to the 7-operator gel shift binding data, holding B/U and F fixed (Supplementary Methods). The fraction of DNA remaining unbound in the Apl 7 gel shift (Fig. 2) was quantified, the number of Apl sites in the model was set to 7, and the model modified by removing all species involving RNAP. This gave association constants B=79,700 M^−1^ and U=17,300 M^−1^ and a good match to the Apl 7 data (Fig. 6F).

**Figure 6.**
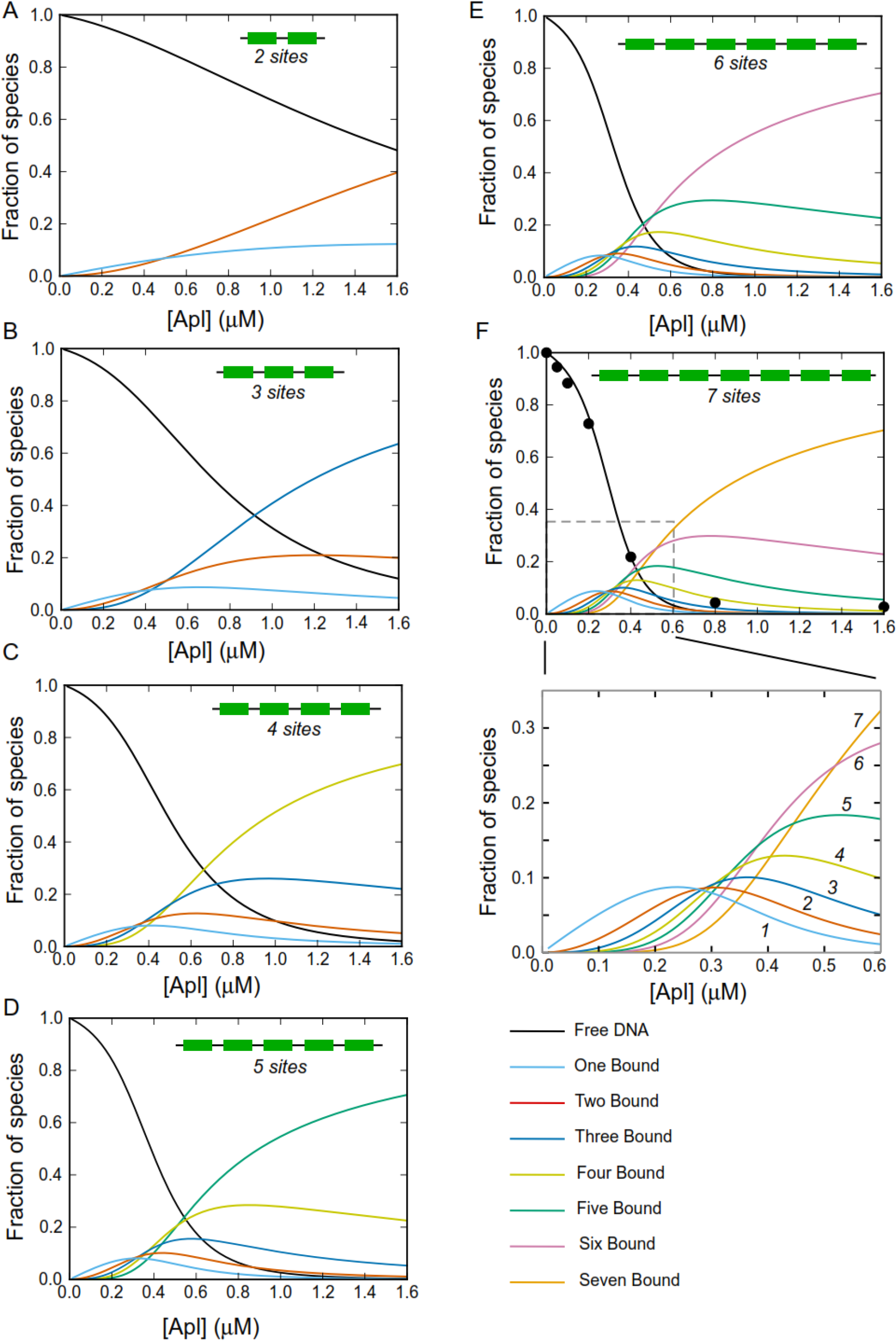
Model prediction of gel shift data. The model was used to predict the proportion of species with different numbers of Apl operators, shown in solid coloured lines, for models with two to seven specific sites (A-E). The seven site gel shift data that was used to calibrate the Apl concentration range is shown with black circles, while panel F shows an enlargement of the concentration regime for the Apl7 model, where several species of different stoichiometries are predicted to co-exist.

These *in vitro* derived parameters were then used to predict binding patterns as a function of Apl concentration for each of the other fragments used in gel shifts (Fig. 2), each time adjusting the model for the numbers of operators and the presence (where applicable) of scrambled sites. The model allows us to calculate the relative proportion of all the possible states, and hence calculate the proportion of each Apl binding stoichiometry. The results, over a range of Apl concentrations, are plotted in Fig. 6. The first point to note is that, as expected, the apparent K_D_, determined as the Apl concentration which gives 50% of DNA unbound, decreases as the number of specific sites increases. This is mirrored in the gel shifts with intact operators, where the apparent K_D_ decreases from ∼ 800 nM for the three operator fragment to ∼ 275 nM for the seven operator fragment. Although the model does predict some weak binding to a two site fragment, we did not detect this experimentally in the gel shift. It is likely that there is rapid dissociation of weakly bound complexes during the course of the gel shift experiment. The simulations also show that binding is much stronger when there are adjacent specific sites. Although the 101, 1101 and 10101 fragments are predicted by the model to bind Apl at high Apl concentrations, the overall affinity was very much weakened (Fig 7 A-F). While our model assumes for simplicity a constant cooperativity value (F) between adjacently bound Apl monomers, it is possible that cooperativity between a specifically bound Apl and an Apl bound to a scrambled site is weaker.

**Figure 7.**
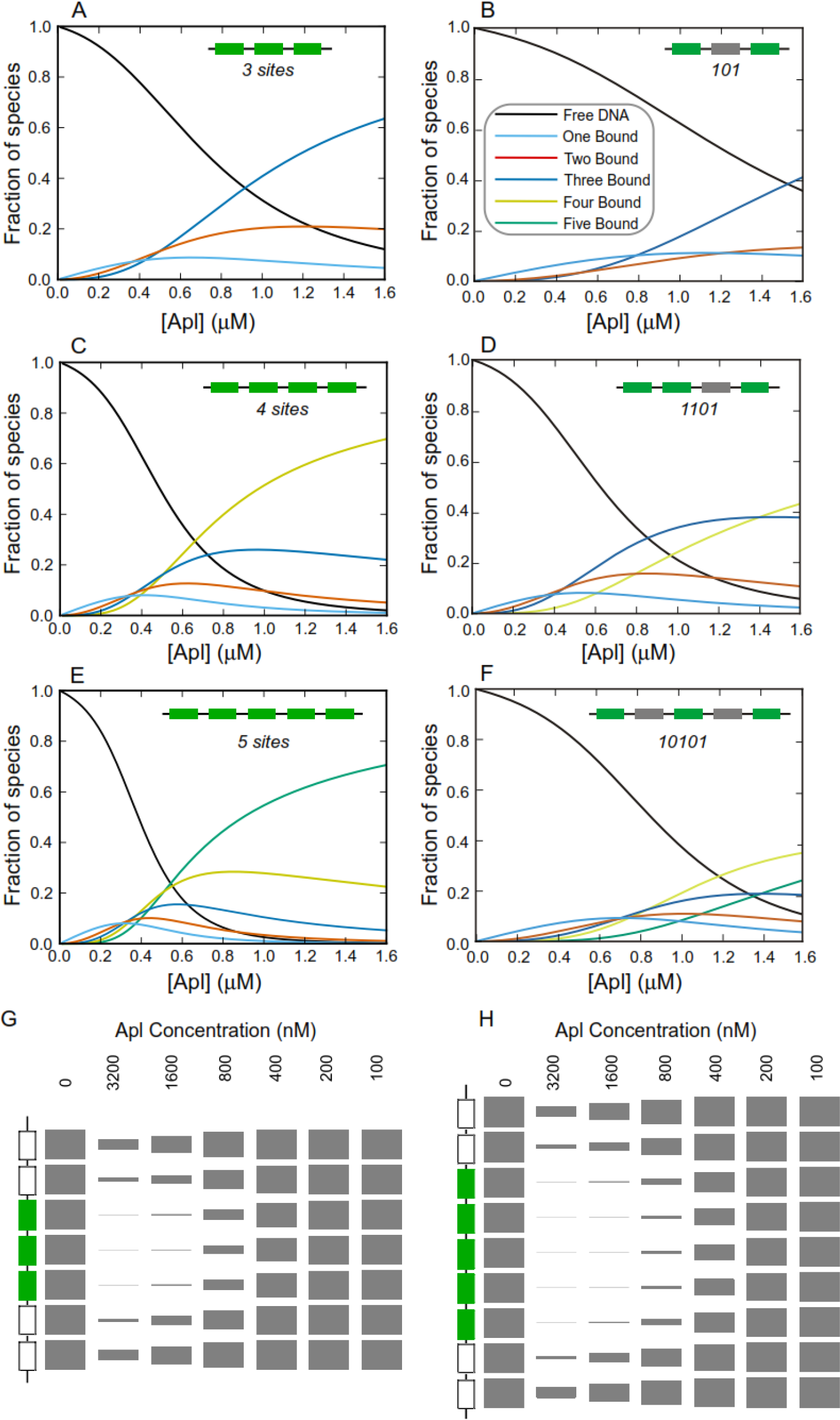
Comparison of the predicted distribution of binding stoichiometries for three (A, B), four (C, D) or five (E, F) operators, with either consecutive wild type (green boxes) (A, C, E) or with intervening scrambled operators (grey boxes) (B, D, F). Simulation of DNase footprinting data for (G) three and (H) five operators, respectively, reproduces the observed spreading of Apl binding into flanking regions.

Thus, the model reproduced the *in vitro* binding data reasonably well. In addition, the fitting enables the affinity of Apl to a single site to be calculated. The binding of Apl to a single operator is very weak, with an estimated K_D_ of 12.5 μM. As a result of the moderate level of cooperativity between bound Apl monomers, the modelling also predicts that at intermediate Apl concentrations (∼0.2-0.7 μM), where all operators are not occupied, significant fractions of species with different numbers of bound Apl monomers should be present (Fig. 6). Indeed, consistent with the predicted distribution of species, two retarded bands were seen for fragments containing three, four or five operators, and more than three bands were apparent with the 6- and 7-site fragments. Uncertainty in how the various Apl-DNA complexes migrate in the gel means it is not possible to assign bands to specific species.

We also used the model to simulate Apl spreading in the DNase I footprints. Two non-specific sites were placed on each side of three and five specific operators, and the probability for each of the sites to be occupied was calculated over a range of Apl concentrations. The result is depicted in ‘footprint form’, showing the expected spreading and the lack of a clear boundary between the specific and non-specific sites (Fig. 7 G-H), comparable to experimental observation.

### Targeted mutations in Apl to reduce binding

Bioinformatic analysis has classified the tyrosine integrase associated RDFs into subgroups based on different predicted DNA binding motifs. The Xis and P2 Cox families are predicted (Nilsson *et al.*, 2011), and in some cases shown experimentally (Abbani *et al.*, 2007)(Berntsson *et al.*, 2014)(Singh *et al.*, 2014), to bind DNA through a winged-helix motif. The Apl-related RDFs on the other hand are strongly predicted (Dodd and Egan, 1990)(Nilsson *et al.*, 2011) but not yet shown, to bind DNA via a conventional helix-turn-helix (HTH) motif.

Structures of tyrosine integrase associated RDFs, where available, suggest that cooperativity in binding is mediated by the interaction of the C-terminal tail of one monomer with an adjacent DNA-bound monomer, in a head to tail manner (Abbani *et al.*, 2007)(Berntsson *et al.*, 2014)(Singh *et al.*, 2014). We asked whether Apl might bind similarly, by mutating Apl and assessing binding activity *in vivo* by repression of a pR-lacZ reporter (Fig 8).

**Figure 8.**
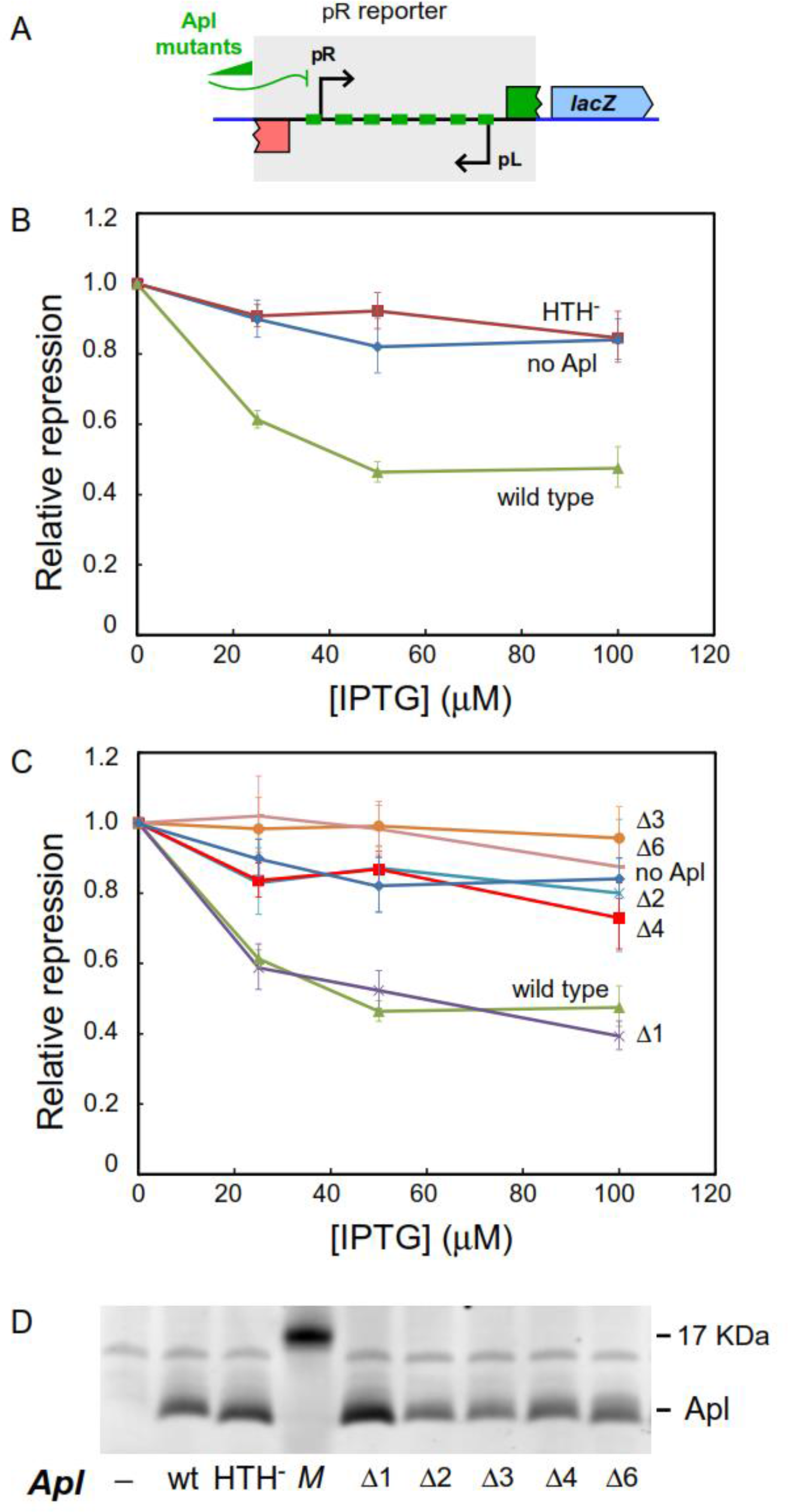
Mutation of the predicted helix-turn-helix or small C-terminal deletions eliminate Apl activity. A. A pR-lacZ reporter was used to measure the ability of Apl mutants to repress pR activity. B. Mutations at key residues in the Apl HTH (E28R/R29E/R33Q) eliminates the ability of Apl to repress the pR-lacZ reporter. C. Apl C-terminal deletions of 2-6 amino acids dramatically reduce pR repression. A single amino acid deletion had little effect of repression. D. Western blots (D) of extracts from the same strains, grown in the presence of 100uM IPTG, showed that Apl expression levels were similar for each of the mutants. M = MW marker.

To test whether the predicted HTH motif is required for binding, we mutated three key residues in the HTH to other residues commonly found at those positions, to give Apl (E28R/R29E/R33Q) (Supplementary Fig. 5). These changes resulted in complete loss of pR-lacZ repression (Fig. 8). To test the contribution of the Apl C-terminal residues to cooperativity, we made a series of single amino acid deletions at the C-terminus (Supplementary Fig. 5). While Apl with a one amino acid deletion retained complete activity, deletions of two or more amino acids resulted in almost complete loss of pR repression. Western blots of whole cell extracts, normalised for OD_600nm_ to ensure equal loading, showed that Apl was produced at near wild type levels for each of the mutants (Fig. 8C). While we cannot rule out that the deletions may somehow more directly affect binding, these results are consistent with the C-terminus playing a role in cooperative interactions between monomers.

## DISCUSSION

### Apl’s DNA binding mode

DNase I footprints of Apl at its binding sites at pR-pL and at *att*P are indicative of Apl binding on the inside face of bent DNA (Dodd, Reed and Egan, 1993). Our results and previous studies establish a number of features of Apl DNA binding, many or all of which are shared with other well-studied RDFs (Saha, Haggard-Ljungquist and Nordstrom, 1989)(Yu and Haggard-Ljungquist, 1993)(Esposito and John J Scocca, 1997)(Lewis and Hatfull, 2001)(Calì *et al.*, 2004)(Abbani *et al.*, 2007)(Mattis, Gumport and Gardner, 2008)(Singh *et al.*, 2014): (1) the operators are arranged as direct repeats, spaced roughly one turn of the DNA helix apart, presumably with one monomer bound per operator; (2) binding to a single specific operator is weak; (3) binding to adjacent operators is highly cooperative; (4) binding causes DNA bending; and (5) the difference in affinity for specific and non-specific sites is small, presumably reflecting flexible sequence recognition. The presence of these features in Apl supports the idea that this basic mode of DNA binding is universal in this group of proteins (Abbani *et al.*, 2007)(Mattis, Gumport and Gardner, 2008).

Taking advantage of Apl’s activity as a transcriptional repressor and by using operator mutants, we were able to generate *in vivo* data that enabled model-based extraction of estimates for the basic biochemical parameters for Apl binding. A simple model that uses two DNA binding affinities for Apl (specific and non-specific) and a single parameter for cooperation between adjacent monomers was able to reproduce the *in vivo* and *in vitro* binding data reasonably well. Monomer binding to a single site was estimated to have K_D_s of ∼12.5 μM (ΔG of −6.9 kcal/mol) for a specific operator, and ∼58 μM (ΔG of −6.0 kcal/mol) for a non-specific site. The fitted value for cooperativity, F ∼50 is equivalent to a ΔG of −2.4 kcal/mol. Although these values should be regarded with caution, they compare reasonably with estimates for other DNA-binding proteins. In gel-shift experiments, a K_D_ of 1 μM was estimated for P22 Xis binding to a DNA with a single operator (Mattis, Gumport and Gardner, 2008), and weak but detectable binding to single operators at nM concentrations was seen for λ Xis, HP1 Cox and Gifsy-1 Xis (Yin, Bushman and Landy, 1985)(Esposito and Scocca, 1997)(Flanigan and Gardner, 2007) suggesting single site affinities that are substantially higher than Apl. Thus, it seems that Apl’s affinity for its specific sites is relatively low, hence more binding sites may be needed in order to compensate. The 12.5 μM K_D_ is an average over all the Apl operators at pR-pL, but as the consensus is not strictly conserved, some operators may have higher affinity than others.

The relative levels of intermediate species seen in gel shifts with multiple operator DNA, and the coordinate occupation of multiple adjacent sites in DNAseI footprinting experiments reflects a balance between binding affinity and cooperativity. Our model allowed us to derive a quantitative measure of cooperativity, a parameter which is not available for other RDFs. Apl’s cooperativity factor of ∼50 is comparable to F = 60-130 between λ CI dimers (= −2.5 to −3 kcal/mol; (Johnson, Meyer and Ptashne, 1979), but less than the ∼2000 (−4.7 kcal/mol) for HK022 repressor dimers (Carlson and Little, 1993).

Hence, the derived values for both binding affinity and cooperativity are comparable to other regulators of expression, and both affinity and cooperativity can be tuned for the desired regulatory outcomes.

### The DNA binding mode and RDF function

The conservation of DNA binding mode among RDFs suggests that it is particularly suited to allow RDFs to foster a specific spatial arrangement of the DNA flanking their binding sites in order to create appropriate DNA substrates for binding of the integrase and other recombination proteins.

Strong cooperativity between adjacent RDFs is likely to be needed to impart a static bend and a stiffening of the DNA to fix an optimal recombination structure. Each additional binding site provides an ‘architectural increment’ to this structure. Thus, a monomer/one-DNA-turn binding unit for an RDF may be an advantage over a typical dimer/two-DNA-turn binding unit because it allows for smaller architectural increments in the evolutionary construction of *att* sites. The ability to spread into adjacent sites provides an ‘RDF concentration window’ for the creation of a particular DNA structure, since one less or more RDF in the chain is likely to significantly change the overall DNA arrangement. How the length of the RDF chain responds to RDF concentration can be readily tuned by alterations in the DNA sequence to foster or hinder the addition of the next monomer in the chain or by alteration in the cooperativity between RDFs. Spreading may also position the RDF where it can make favourable or competitive contacts with other recombination proteins, providing further concentration-dependent regulation of recombination. Mattis et al. (Mattis, Gumport and Gardner, 2008) proposed that occupation of non-specific sites that overlap Int binding sites may cause high concentrations of P22 Xis to inhibit reintegration after excision.

### The DNA binding mode and transcriptional regulation

Although the DNA binding properties of Apl and other RDFs seem well suited to their recombinational role, two features can also be used to provide effective and unusual transcriptional regulation. The first is highly cooperative binding, a feature that is common among transcription factors and which is used to generate sharp transitions between promoter activity and inactivity in response to small changes in regulator concentration (Hope, Rebay and Reinitz, 2017). The second is spreading. Spreading from specific sites into non-specific sites is not often used in transcriptional control. One example is the ParB family of proteins, which mediate chromosome partitioning in various replicons. These are dimeric HTH proteins that bind to specific ‘centromere’ sites but also are capable of spreading kilobase distances into adjacent DNA (Sanchez *et al.*, 2015). This spreading is able to silence adjacent genes and may play a role in the partitioning process. Similar spreading has been observed for the *Drosophila* transcription factor Yan and its human homolog TEL/ETV6 (Hope, Rebay and Reinitz, 2017). The spreading of an RDF from its specific operators at *att* could in theory allow repression of the promoter for the recombinase gene when this is located adjacent to *att* but somewhat distant from the primary RDF binding sites. However, in P4 and KplE1, specific RDF sites at *att* overlap the integrase promoter (Calì *et al.*, 2004)(Piazolla *et al.*, 2006)(Panis, Méjean and Ansaldi, 2007), so spreading is not required, at least in these cases.

A spreading mechanism seems to be used in control of the lytic and lysogenic promoters of many P2 related bacteriophages. In these phages, the lytic and lysogenic promoters are arranged face-to-face and the primary operators for the immunity repressor lie over the lytic promoter, while the Apl/Cox operators are distinct from these and tend to lie over the lysogenic promoter or between the transcriptional start sites (Nilsson *et al.*, 2011). In all tested cases, the Apl/Cox proteins repress the lysogenic promoter (Saha, Haggard-Ljungquist and Nordstrom, 1989) (Reed *et al.*, 1997)(Esposito, Wilson and Scocca, 1997), and in some cases also the lytic promoter. It is not clear to what degree the ability of Apl to spread into non-specific sites is important for its regulation of pL and pR, since specific sites overlap both promoters. If binding to non-specific sites is removed in the model, then repression is weakened, but only slightly (repression can be restored by small increases in B or F). However, in P2, Cox repression of the pe early lytic promoter at high concentrations may be due to its spreading from its specific sites over the lysogenic promoter into non-specific sequences at pe. The face-to-face arrangement of the lytic and lysogenic promoters in P2-like phages provides sequence between the promoters that can be used to specify Apl/Cox binding and to set the required balance of repression by adjustment of spreading, without compromising the sequences for RNAP recognition or for immunity repressor binding. In contrast, in the lambdoid phages, the arrangement is more compact, with the lytic and lysogenic promoters back-to-back and the immunity repressor and Cro protein sharing the same operators. Thus, those features that make Apl and the other Cox proteins able to function as RDFs also adapt them well for their roles as lytic regulators of the face-to-face lytic and lysogenic promoters.

## Acknowledgements

This work was supported by funding from the Australian Research Council (DP110100824 and DP150103009).

