## Supplementary materials for "A quantitative binding model for the Apl protein, the dual purpose recombination-directionality factor and lysis-lysogeny regulator of bacteriophage 186"

**SUPPLEMENTARY MATERIAL**

### 1. Supplementary Methods

#### Statistical Mechanical Modelling of Apl binding

#### *In vivo* modelling

Initial modelling aimed to fit the *in vivo* LacZ reporter data, using the seven specific Apl sites between the pR and pL promoters and two non-specific flanking sites on either side of the specific sites, making a total of 11 sites. All possible Apl bound states could then be described as an 11 digit binary number, with a 1 if Apl is bound that site or a 0 if Apl is not, resulting in $2^{11}$ (=2048) different Apl bound states.

The weight for each of these states was stored in an array $2^{11}$ long. Each state is given the initial weight of 1, for each non-specific site occupied each weight multiplied by cU, where U is the non-specific binding parameter, and c is the concentration, and multiplied by c(B+U), where B is the specific binding parameter. For all states, where any site, i, and neighbouring site, i+1, are bound by Apl, the weight is multiplied by the cooperatively parameter, F.

Binding at the first two non-specific and the first two specific sites (i.e. sites 1-4) was considered to compete with RNA polymerase (RNAP) at the pR promoter, and the last two non-specific and last specific sites (ie sites 9-11) considered to compete at the pL promoter, where RNAP covers at least -45 to +10 of a promoter region. Additional states were defined where RNAP could bind to pR when sites 1-4 were not bound by Apl and multiplied by a pR binding parameter, R. The same was done with pL with a pL binding parameter, L. This makes the reasonable assumption that RNAP is at a fixed cellular concentration. Transcription, hence LacZ gene expression from either the pR or pL promoter is considered to occur from all states where RNAP can bind the promoter, hence relative expression is proportional to the probability of RNAP being at the promoter. The probability of RNAP occupying the promoter was determined as the sum of all states where RNAP is at the promoter, divided by the sum of all possible states. To relate this to relative expression Lac Z data, this is normalised by a constant that is related to pR and pL, such that at an Apl concentration of 0, relative expression is 1.

#### Fitting *in vivo* repression data

Four constructs were assayed for Apl binding, each with a different number of specific sites from the seven present in wild type. The arrangements were:

- pR 00111111100 pL
- pR 00110101100 pL
- pR 00110001100 pL
- pL 00111111100 (-pR)

where 0 is a non-specific site, 1 is a specific site, pR represents a pR promoter, pL represents a pL promoter, (-pR) represents a mutant inactive pR promoter and the promoter on the left indicates the promoter from which the LacZ reporter is expressed.

Each different arrangement of Apl sites was modelled as described above, and the relative expression curves were globally fit.

#### Data fitting

The model was fitted to experimental data derived from LacZ assays. In these assays, a LacZ reporter gene was expressed from either a pR or a pL promoter integrated into the bacterial chromosome. Apl was supplied from an IPTG inducible promoter on a separate plasmid, pZE15Apl. Each LacZ assay was repeated on 8-12 biological replicates, and relative expression was determined for each IPTG concentration by dividing by the value at 0 μM IPTG for each replicate. Values for all replicates were used in data fitting, with the error between the model and data for each dataset divided by the total number of points in that data set.

The minimum error was found using a combined random Monte Carlo and linear search method. Random parameter guesses were made, and the best guesses were used as starting guesses for linear optimisation. The advantage of this method is that it avoids local minima, while searching a wide range of parameter values. Initially, 1000 random guesses were taken, and the error minimised with the **fmincon** function of MATLAB. This was performed 100 times. The error as a function of each parameter value was examined, the bounds of the random guesses updated and the minimisation was performed an additional 200 times. Values for both rounds were pooled together, and each parameter plotted against the error to assess convergence. Plotting each parameter value against the error for the 300 rounds of minimisation clearly showed convergence to the lowest error (Supplementary Fig. 3). The twenty fits with the lowest errors were then averaged and the standard deviation determined, resulting in B=4.55 (±0.53) x$10^{-5}$ Apl expression units, U=0.99 (±0.09) x$10^{-5}$ Apl expression units, F=50.7 (±7.9), pR=6.88 (±1.09), pL= 0.29 (±0.15), with an error of 0.37.

The parameter values with the lowest 100 errors were also plotted to investigate parameter correlations (Supplementary Fig. 4). The effect of parameter variation was also examined, where for clarity the species distribution a three site operator sequence was simulated, varying one parameter at a time by +/- 1 standard deviation.

#### Fitting of *in vitro* data

Fitting of *in vitro* binding data was used as an independent test of the model.

Firstly, this was done using gel shift data. Gel shift data was quantified with Imagequant software. The total volume of each lane and the volume of the free DNA of each lane was quantified separately, corresponding to total loaded DNA and unbound DNA. The fraction of free DNA was determined as the amount of unbound DNA divided by the total loaded DNA. The amount of free DNA decreased with increasing Apl concentration, down to a limit which corresponds to a small proportion of DNA that is not annealed correctly and hence is unable to bind Apl. The fraction of free DNA was then normalised to 1 when the Apl concentration was 0, and normalised to 0 when free DNA plateaus. This was done for the seven site gel shift data, for concentrations 0, 50, 100, 200, 400, 800 and 1600 nM of Apl.

As gel shift data was used as an independent measure of the predictive power of the model, the mean of the parameter values for B, U, and F from the *in vivo* fitting, given above, were used. The promoter parameters pR and pL were not used as RNA polymerase is not present in the gel shift assays. To convert B and U into nM units from Lac units, these were multiplied by a calibration factor, d, which was determined by fitting the seven site gel shift data for free DNA using linear optimisation, resulting in a fit value of d=1.75. The resultant fit to the data was excellent, considering only the concentration was rescaled (Figure 6) resulting in B=79,700 $M^{-1}$ and U=17,300 $M^{-1}$.

These same parameter values were used to calculate the distribution of binding stoichiometries as a function of Apl concentration for all fragments used in gel shift assays (Fig 6).

### 2. Supplementary Figures

Oligonucleotides for Apl binding

**A.** pR-pL

Apl 2
TTGA**TGGCAA**GTGT**TGGCAA**ACAG
AACT**ACCGTT**CACA**ACCGTT**TGTC

Apl 3
TTGA**TGGCAA**GTGT**TGGCAA**ACAG**AGTCAA**ATCA
AACT**ACCGTT**CACA**ACCGTT**TGTC**TCAGTT**TAGT

Apl 4
TTGA**TGGCAA**GTGT**TGGCAA**ACAG**AGTCAA**ATCAA**TTGCAA**ACTT
AACT**ACCGTT**CACA**ACCGTT**TGTC**TCAGTT**TAGTT**AACGTT**TGAA

Apl 5
AACCC**ACGCAA**TTGA**TGGCAA**GTGT**TGGCAA**ACAG**AGTCAA**ATCAA**TTGCAA**ACTT
TTGGG**TGCGTT**AACT**ACCGTT**CACA**ACCGTT**TGTC**TCAGTT**TAGTT**AACGTT**TGAA

Apl 6
TATTT**TGGCTA**AACCC**ACGCAA**TTGA**TGGCAA**GTGT**TGGCAA**ACAG**AGTCAA**ATCAA**TTGCAA**ACTT
ATAAA**ACCGAT**TTGGG**TGCGTT**AACT**ACCGTT**CACA**ACCGTT**TGTC**TCAGTT**TAGTT**AACGTT**TGAA

Apl 7
TATTT**TGGCTA**AACCC**ACGCAA**TTGA**TGGCAA**GTGT**TGGCAA**ACAG**AGTCAA**ATCAA**TTGCAA**ACTT**TGGCTA**ATAGG
ATAAA**ACCGAT**TTGGG**TGCGTT**AACT**ACCGTT**CACA**ACCGTT**TGTC**TCAGTT**TAGTT**AACGTT**TGAA**ACCGAT**TATCC

Apl 101
TGA**TGGCAA**GTGT***CTTAGG***ACAG**AGTCAA**ATCA
ACT**ACCGTT**CACA***GAATCC***TGTC**TCAGTT**TAGT

Apl 1101
AACCC**ACGCAA**TTGA**TGGCAA**GTGT***CTTAGG***ACAG**AGTCAA**ATCA
TTGGG**TGCGTT**AACT**ACCGTT**CACA***GAATCC***TGTC**TCAGTT**TAGT

Apl 10101
AACCC**ACGCAA**TTGA***CTTAGG***GTGT**TGGCAA**ACAG***CTTAGG***ATCAA**TTGCAA**ACTT
TTGGG**TGCGTT**AACT***GAATCC***CACA**ACCGTT**TGTC***GAATCC***TAGTT**AACGTT**TGAA

Apl 1110111
TATTT**TGGCTA**AACCC**ACGCAA**TTGA**TGGCAA**GTGT***CTTAGG***ACAG**AGTCAA**ATCAA**TTGCAA**ACTT**TGGCTA**ATAGG
ATAAA**ACCGAT**TTGGG**TGCGTT**AACT**ACCGTT**CACA***GAATCC***TGTC**TCAGTT**TAGTT**AACGTT**TGAA**ACCGAT**TATCC

Apl 1101011
TATTT**TGGCTA**AACCC**ACGCAA**TTGA***CTTAGG***GTGT**TGGCAA**ACAG***CTTAGG***ATCAA**TTGCAA**ACTT**TGGCTA**ATAGG
ATAAA**ACCGAT**TTGGG**TGCGTT**AACT***GAATCC***CACA**ACCGTT**TGTC***GAATCC***TAGTT**AACGTT**TGAA**ACCGAT**TATCC

**B.** *att*P

*att*P 5
ATCA**TTGACA**TAGA**TTGCCA**TTGT**TTGCCA**ATAA**TTACCT**ATTA**TTCGCC**ATTTC
TAGT**AACTGT**ATCT**AACGGT**AACA**AACGGT**TATT**AATGGA**TAAT**AAGCGG**TAAAG

**Supplementary Figure 1.** Apl binding site sequences used to assess Apl binding at the pR-pL region (A) or *att*P region (B). Double stranded sequences containing Apl binding sites (bold) or scrambled sites (bold, italics) are shown. Underlined sequences indicate the oligonucleotides used to generate the double stranded sequences.

A.

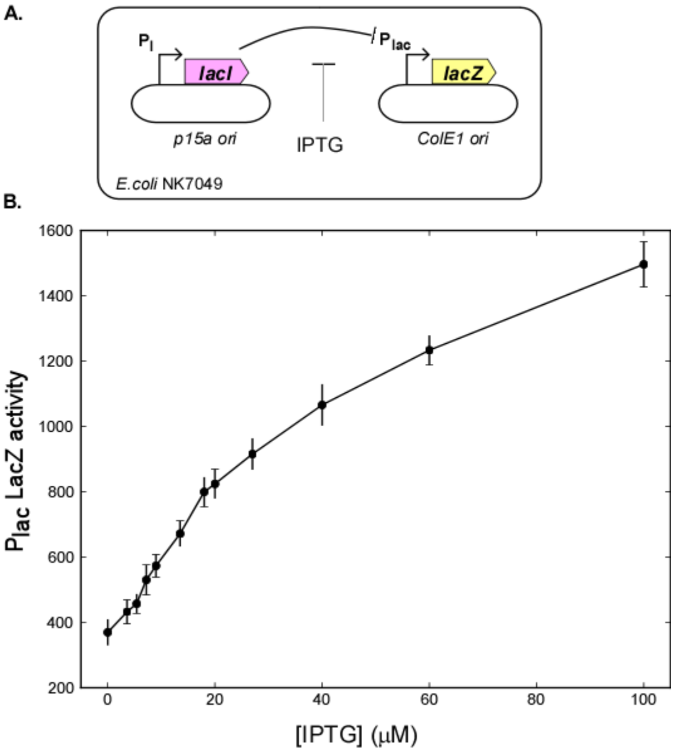


**Supplementary Figure 2.** Relationship between IPTG concentration and pLac activity used to estimate relative expression levels of Apl *in vivo*. The *in vivo* Apl expression system (Fig. 5) used an identical plasmid set up to that shown here, where the *lacZ* gene has replaced the *apl* gene (A). Thus, relative Apl expression units at different IPTG concentrations can be obtained by using the LacZ vs IPTG plot shown here (B). Error bars represent 95% confidence intervals.

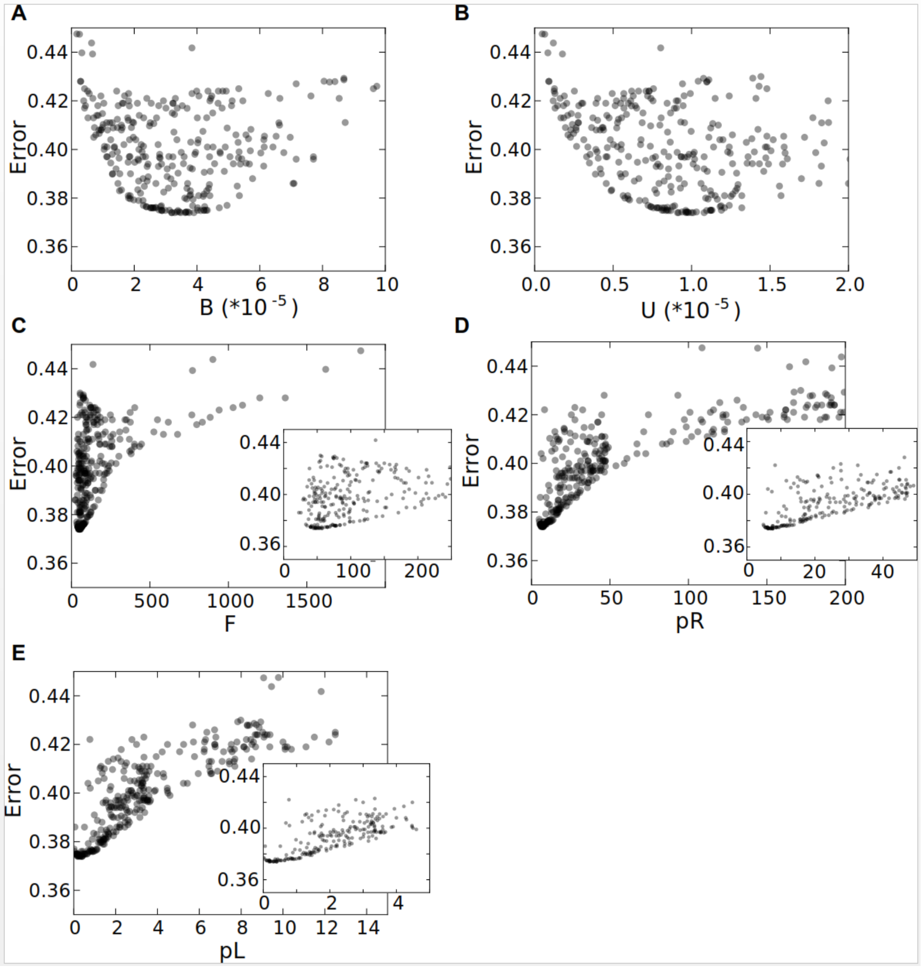


**Supplementary Figure 3.** Optimisation of best fit parameters.

Error terms in the fitting are plotted against the values of the parameters for specific Apl binding (panel A), non-specific binding (panel B), cooperativity (panel C), pR (panel D) and pL (panel E). The plots show that the fitting procedure converges to minimise the error term. Insets in panels C-E show more detail.

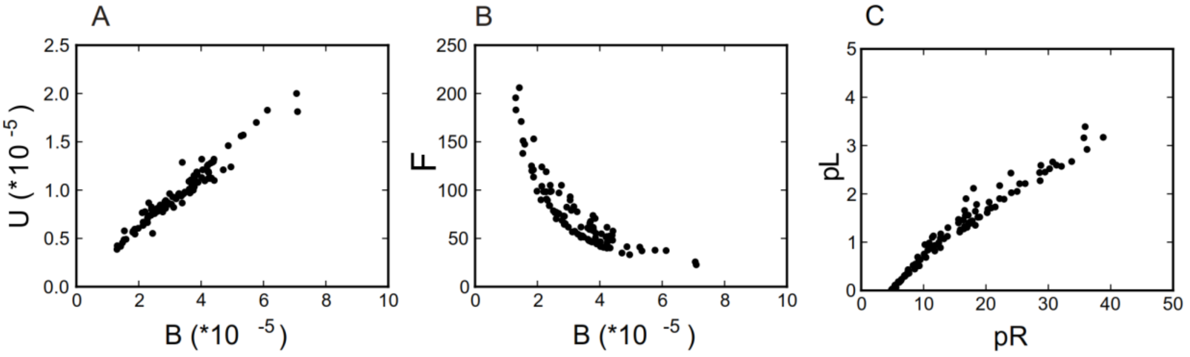


**Supplementary Figure 4.** Correlation between fitted parameters.

Parameter correlations for the best 100 fits of the model to the lacZ data of Figure 5 are shown here.
(Panel A) Non-specific binding (U) increases linearly with specific binding strength (B).
(Panel B) Cooperativity (F) decreases as specific binding strength (B) increases.
(Panel C) Promoter strengths for pR increased linearly with pL strength.

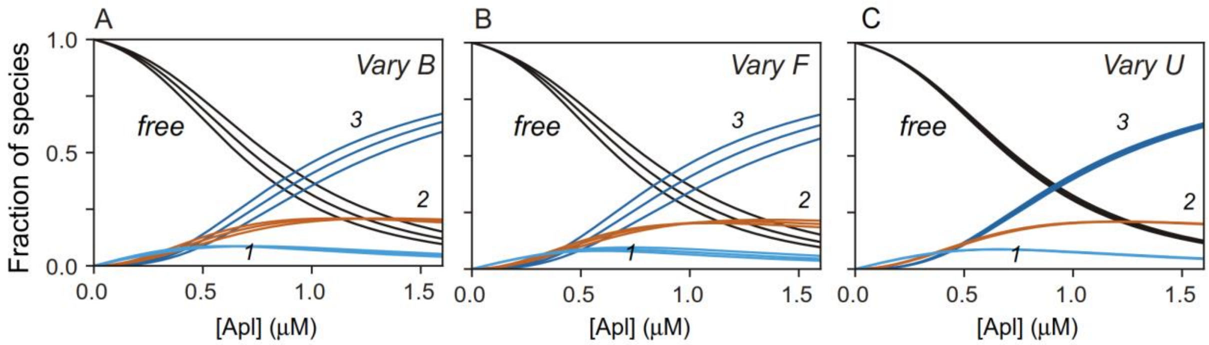


**Supplementary Figure 5.** Effect of parameter variation on species distributions.

The range of species distributions are plotted for the case of three consecutive Apl operators. For each plot, two of the three parameters are held fixed at their mean values and the third parameter varied higher and lower by one standard deviation. The standard deviation of the parameters was obtained from the best 100 fits to the data shown in Figure 5. As expected, variation in U (panel C) has no impact when the DNA contains only specific sites.

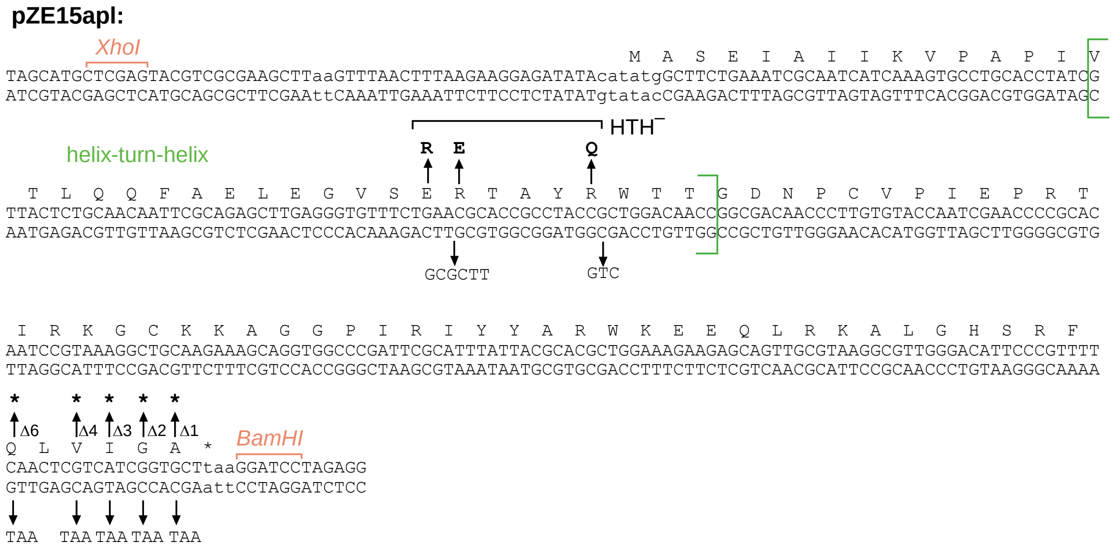


**Supplementary Figure 6:** Details of Apl mutants.

The pZE15-Apl plasmid (*Xho*I to *Bam*HI region shown here) was mutated as shown to produce HTH$^{-}$ and C-terminal deletion mutants. The predicted helix-turn-helix motif is indicated within the green brackets.
